# The dynamics of cortical GABA in human motor learning

**DOI:** 10.1101/341503

**Authors:** James Kolasinski, Emily L. Hinson, Amir P. Divanbeighi Zand, Assen Rizov, Uzay E. Emir, Charlotte J. Stagg

**Affiliations:** Oxford Centre for fMRI of the Brain, Wellcome Centre for Integrative Neuroimaging, Nuffield Department of Clinical Neurosciences, University of Oxford, UK, OX3 9DU; Cardiff University Brain Research Imaging Centre, School of Psychology, Cardiff University, UK, CF24 4HQ; Oxford Centre for Human Brain Activity, Wellcome Centre for Integrative Neuroimaging, Department of Psychiatry, University of Oxford, Oxford, UK, OX3 7JX; Perdue University School of Health Sciences, 550 Stadium Mall Drive, West Lafayette, IN 47907, USA

## Abstract

The ability to learn novel motor skills is both a central part of our daily lives and can provide a model for rehabilitation after a stroke. However, there are still fundamental gaps in our understanding of the physiological mechanisms that underpin human motor plasticity. The acquisition of new motor skills is dependent on changes in local circuitry within the primary motor cortex (M1). This reorganisation has been hypothesised to be facilitated by a decrease in local inhibition via modulation of the neurotransmitter GABA, but this link has not been conclusively demonstrated in humans. Here, we used 7T MR Spectroscopy to investigate the dynamics of GABA concentrations in human M1 during the learning of an explicit, serial reaction time task. We observed a significant reduction in GABA concentration during motor learning that was not seen in an equivalent motor task lacking a learnable sequence, nor during a passive resting task of the same duration. No change in glutamate was observed in any group. Furthermore, baseline M1 GABA was strongly predictive of the degree of subsequent learning, such that greater inhibition was associated with poorer subsequent learning. This result suggests that higher levels of cortical inhibition may present a barrier that must be surmounted in order achieve an increase in M1 excitability, and hence encoding of a new motor skill. These results provide strong support for the mechanistic role of GABAergic inhibition in motor plasticity, raising questions regarding the link between population variability in motor learning and GABA metabolism in the brain.

**Funding information:** J.K.:Wellcome Trust Sir Henry Wellcome Postdoctoral Fellowship (204696/Z/16/Z). C.J.S.: Wellcome Trust/Royal Society Henry Dale Fellowships (102584/Z/13/Z).

## 1 INTRODUCTION

Motor learning describes the process by which we change and adapt in our interactions with the external world (Dayan and Cohen, 2011). The ability to acquire new motor skills has been strongly associated with plastic changes both the in structure and functional organisation of the primary motor cortex (M1) (Dayan and Cohen, 2011; Sampaio-Baptista et al., 2018). Specifically, evidence from both human and non-human primate studies suggests that repeated practice of a motor skill is associated with changes in the topographic organisation of the region (Nudo et al., 1996; Karni et al., 1998). Further, the learning of fine motor skills has been associated with synaptogenesis in M1 (Kleim et al., 2002), as well as changes in the myelination of the underlying white matter (Sampaio-Baptista et al., 2013). Understanding the physiological processes that drive the observed structural and functional changes in M1 to support motor learning are necessary for the development of therapeutic approaches to promote adaptive plasticity after brain injuries, such as stroke, via facilitation of the re-learning of motor skills compromised by brain pathology.

There is a strong body of evidence to suggest that a reduction in cortical inhibitory tone is critical for the induction of M1 plasticity (Bachtiar and Stagg, 2014; Peters et al., 2017). Specifically, a reduction in *γ*-Aminobutyric acid (GABA)-ergic signalling appears crucial to the induction of LTP-like plasticity in M1 (Trepel and Racine, 2000; Castro-Alamancos et al., 1995), potentially by unmasking or potentiating latent pre-existing horizontal connections in the cortex (Huntley, 1997). In addition, particularly compelling evidence for the role of GABAergic disinhibition in promoting M1 plasticity is provided by recent work using in-vivo two-photon imaging in mouse M1. Learning resulted in a significant reduction in axonal boutons observed on somatostatin-expressing inhibitory neurons (Chen et al., 2015). Optogenetically manipulating activity in this inhibitory neuronal population during learning disrupted both the observed dendritic structural changes and also affected motor performance.

Changes in human M1 GABAergic activity have been observed during human motor learning using Transcranial Magnetic Stimulation (TMS) (Rosenkranz et al., 2007), though this has not been consistently reported. Using Magnetic Resonance Spectroscopy (MRS), decreases in GABA in response to the non-invasive brain stimulation approach transcranial direct current stimulation (tDCS) have been shown to be predictive of learning a motor task in healthy controls(Stagg et al., 2011a). Evidence for a reduction in cortical GABA has also previously been reported in the context of motor learning over the course of weeks, with correlated changes in the strength of connectivity in the resting state motor network as a whole (Sampaio-Baptista et al., 2015). Lower levels of M1 GABA have also been reported in the chronic stages of recovery after stroke, relative to unaffected individuals; in these patients a reduction in M1 GABA is associated with a favourable clinical response to a therapeutic intervention (Blicher et al., 2015). However, to date, only one study has reported direct evidence for a reduction in MRS-assessed M1 GABA in humans during the learning of a new motor skill (Floyer-Lea, 2006); a result which has proven difficult to replicate.

Here, we take advantage of the increased spatial, temporal, and spectral resolution afforded by acquiring MRS data at ultra-high field (7T) to investigate the changes in M1 GABA in M1 during motor learning (Figure 1). We sought to address the hypothesis that MRS-assessed M1 GABA will decrease during learning of a motor task, and further that GABA concentration early in learning will predict the magnitude of subsequent motor learning (Stagg et al., 2011a).

**FIGURE 1.**
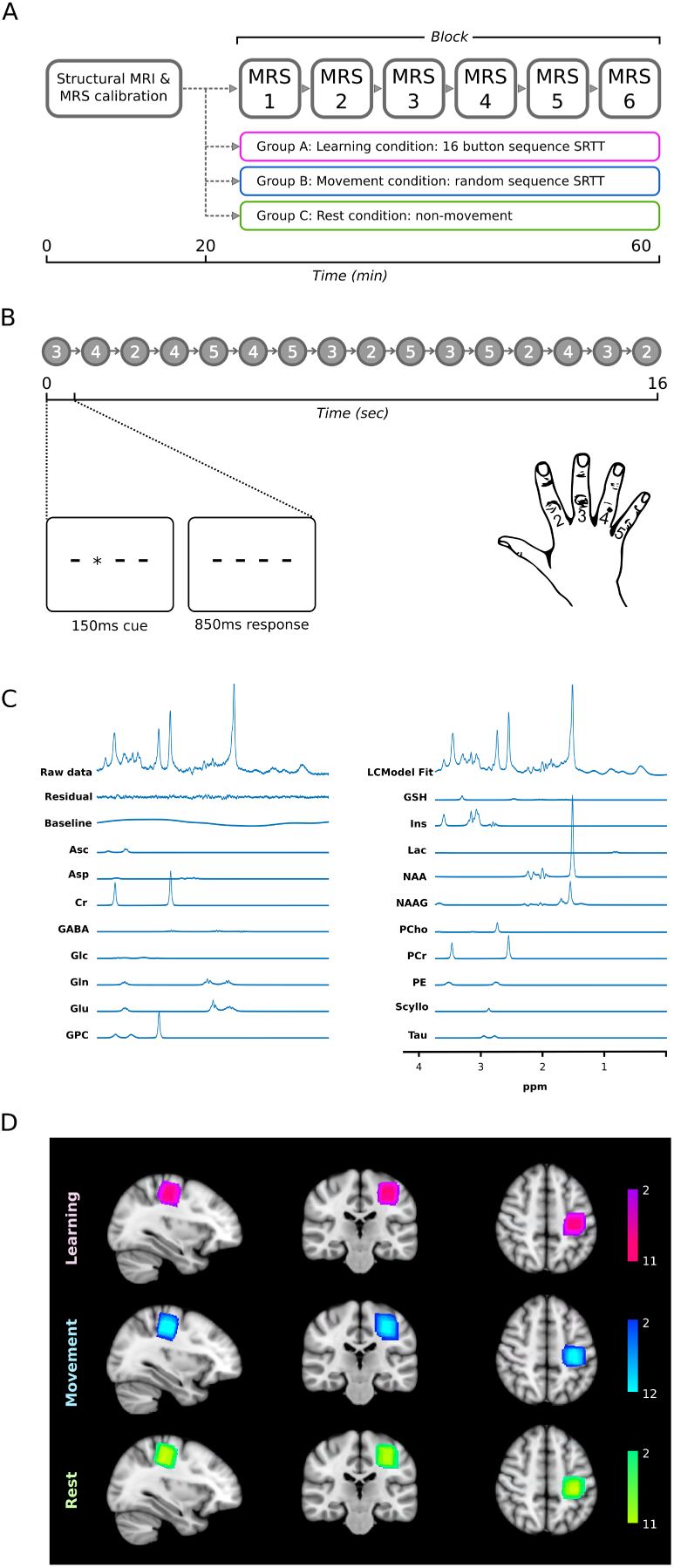
Experimental design and MR spectroscopy data acquisition. MRS data were acquired in six independent blocks during a concurrent task which differed across the three experimental groups (A): the Learning group performed a 16-button press repeating serial reaction time task; the Movement group performed a serial reaction time task without a repeating sequence; the Rest group passively observed a video. The Learning group performed the same 16-button press sequence (B): 3 repeats per epoch (48 seconds), with each epoch separated by a 12 second rest period. MRS data were acquired in each block (64 averages) and analysed using LCModel (Provencher, 2001) (C): representative acquisition from one participant in one M1 block, including model fit). (D): M1 MRS voxels were centred over the left (contralateral) hand knob illustrated in as heatmaps in the three experimental groups: colour bars represent number of participants.

## 2 MATERIALS AND METHODS

### 2.1 Participants

51 healthy participants gave written informed consent to participate in the study (Oxford University Central Research Ethics Committee: MSD-IDREC-C1-2014-100). Participants were right handed according to the Edinburgh Handedness Questionnaire (Oldfield, 1971) and met local safety criteria for scan participation at 7T. Participants were allocated to one of three experimental groups: Learning, Movement, or Rest (Figure 1A). A full breakdown of group allocation is provided in Table 1. No participant was recruited to more than one group. All participants attended one scanning session.

**TABLE 1.**
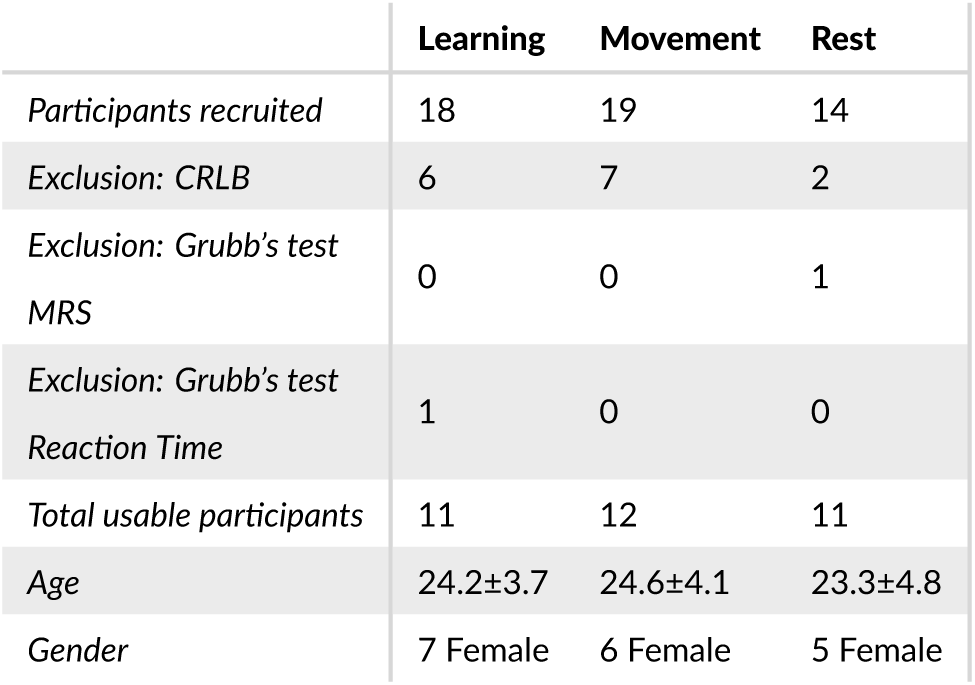
Participant breakdown across experimental groups

|  | Learning | Movement | Rest |
| --- | --- | --- | --- |
| Participants recruited | 18 | 19 | 14 |
| Exclusion: CRLB | 6 | 7 | 2 |
| Exclusion: Grubb's test<br>MRS | 0 | 0 | 1 |
| Exclusion: Grubb's test<br>Reaction Time | 1 | 0 | 0 |
| Total usable participants | 11 | 12 | 11 |
| Age | 24.2±3.7 | 24.6±4.1 | 23.3±4.8 |
| Gender | 7 Female | 6 Female | 5 Female |

### 2.2 MR data acquisition

MRI and MRS data were acquired using a 7 tesla Siemens Magnetom System (Siemens, Erlangen, Germany) with a 32-channel head coil (Nova Medical Inc, Wilmington, MA, USA). A dielectric pad (barium titanate: 0.5 × 11 × 11 cm) was positioned dorsally on the scalp over left central sulcus to increase B1 efficiency in the M1 voxel of interest (VOI) (Lemke et al., 2015). B1 efficiency was imaged using actual flip angle imaging (AFI): FOV 240×240, TR1 6 ms, TR2: 30 ms, TE 2.58 ms, non-selective flip angle 60°, slice thickness 2.5 mm.

To enable placement of the MRS voxel, structural MRI data were acquired with a Magnetization Prepared Rapid Acquisition Gradient Echo (MPRAGE) sequence: TR=2200 ms, TE=2.82 ms, slice thickness 1.0 mm, in-plane resolution 1.0 × 1.0 mm, GRAPPA factor = 2.

MRS data were acquired using a semi-LASER sequence (van de Bank et al., 2015): TR=5000 ms, TE=36 ms, 20×20×20 mm voxel, 64 averages per block, TA=5 mins 20 secs, using VAPOR (VAriable Power RF pulses with Optimized Relaxation delays) water suppression (Tkác et al., 1999). The VOI was manually positioned in the left M1, covering the whole hand knob (Yousry et al., 1997) (Figure 1D) and excluding the dura. MRS data were acquired in six blocks of approximately 5 minutes each (Figure 1A). During the acquisition of the MRS data, participants performed either an explicit sequence learning task, a motor task without a learnable sequence, or watched a video.

### 2.3 MRS task stimuli

Both the Learning and Movement groups were engaged in a visually-cued serial reaction time task (SRTT). Responses were made with the right hand via a four-button button box resting on the participants’ thigh. Visual cues consisted of four horizontal lines displayed on a screen, representing the four buttons (Figure 1B). Each cue consisted of one line being replaced with an asterisk for 150 ms. Participants were instructed to respond by pressing the corresponding button as quickly and accurately as possible, and not to press in anticipation of an upcoming cue. There was an interstimulus interval (ISI) of 850 ms between cues. 48 cues were presented in each epoch, followed by a rest period of 12 seconds. The task repeated throughout MRS acquisition (Figure 1A).

In the Learning group participants were explicitly informed to expect a repeating sequence in the cues (Figure 1B: a 16-item sequence repeated three times per epoch). In the Movement group, participants were told not to expect a sequence; cues were pseudo-randomised to produce different sequences of 48 cues in each epoch, the number of button presses for each finger was matched to the learning task.

In the Rest group, participants watched a 40 minute excerpt from a nature documentary. Between each MRS block, participants were cued to press a button.

### 2.4 MR data analysis

Raw MRS data from each block separately were corrected using the unsuppressed water signal acquired from the same VOI, and were subject to eddy current correction and a zero-order phasing of array coil spectra using in-house scripts. Any residual water signal was removed using Hankel-Lanczos singular value decomposition (Cabanes et al., 2001). LCModel analysis was used to quantify a concentration of neurochemicals within the chemical shift range 0.5 to 4.2 ppm (Provencher, 2001). The exclusion criteria for data were as follows: Cramer-Rao bounds (CRLB) > 50%, water linewidths at full width at half maximum (FWHM) > 15 Hz, or SNR < 40. There was no strong correlation (> ±0.3) between GABA and other metabolites, indicating good spectral separation was achieved.

To quantify the proportion of white matter (WM), grey matter (GM), and cerebrospinal fluid (CSF) in the VOI, FMRIB’s Automated Segmentation Tool (FAST) (Zhang et al., 2001) was applied to the T1-weighted MPRAGE scan. GABA and glutamate peaks were corrected for the proportion of GM in the VOI. Total creatine (including phosphocreatine: tCr) peaks were corrected for the proportion of total brain tissue in the VOI.

MRS data analysis therefore yielded independent quantification of neurochemical concentrations corresponding of each of the six MRS blocks, expressed as a ratio of tCR, for example GABA:tCr and Glu:tCr (Figure 1A).

### 2.5 MRS motor task data analysis

In the Learning and Movement groups, task performance was assessed by quantifying response time (RT) between stimulus presentation and button press response for correct response only. RT data were divided into blocks corresponding to the six independent MRS acquisitions. Median RT were then calculated within each block for each participant and used for subsequent analysis. The summary measure of learning was defined as the difference between the median RT in block 1 and the lowest reaction time across blocks 2-6. This metric aimed to capture the peak learning, which, dependent on the rate of learning, might not necessarily occur in the block 6, particularly in participants who exhibit rapid learning early in the task.

### 2.6 Statistics

Statistical analyses and graphing were conducted using JMP (Version 13.0, SAS Institute, Cary, NC, USA) and Statistics Package for the Social Sciences (SPSS, Version 22.0, IBM Corporation, Armork, NY, USA). To compare changes in RT across the Learning and Movement groups, median RT data for each block were subject to a two-way mixed ANOVA: Between-subjects factor: experimental group (Learning or Movement), Within-subjects factor: time (Block 1-6). To make similar comparisons regarding changes in the MRS concentration of GABA across the three experimental groups, GM-corrected GABA:tCr values were also subject to a two-way mixed ANOVA: Between subjects factor: experimental group (Learning, Movement, or Rest), Within-subjects factor: time (Block 1-6). Significant interactions were followed up with analysis of simple main effects within each experimental group. Correlations were assessed using Pearson’s correlation coefficients (two-tailed). Comparison of correlation statistics was undertaken using Hittner’s test (Hittner et al., 2003). A total of 17 participants were excluded on the basis of pre-defined criteria (Table 1): 15 participants were excluded on the basis of their MRS CRLB exceeding 50%; one participant was excluded from the Rest group due to a statistical outlier in their GABA:tCr values (±2 SD); one participant was excluded from the Learning group due to a statistical outlier in the median RT values (±2 SD). Normality of the remaining data was confirmed using Shapiro-Wilk tests.

## 3 RESULTS

### 3.1 Motor sequence learning is associated with a reduction in M1 GABA concentration

We observed a significant reduction in MRS measures of M1 GABA over time in the Learning group, in line with the significant reduction in RT over time in this group. No change was observed in RT in the Movement group, and no reduction in GABA:tCr was observed in either the Movement group, nor the Rest group.

In the analysis of RT data, a two-way mixed ANOVA revealed a significant interaction between experimental group (Learning or Movement) and time on RT: F_(1.49,105)_ = 10.52 p=0.001, partial *η*^2^ = 0.334 (Greenhouse Geisser corrected), therefore simple main effects were analysed (Figure 2). In the Learning group there was a significant main effect of time on RT: F_(1.41,14.0)_ = 9.33 p=0.005, partial *η*^2^ = 0.483 (Greenhouse Geisser corrected). Post-hoc paired-samples t-tests revealed a significant reduction in RT in the Learning group in blocks 4 and 6 compared with block 1 (p<0.05, Bonferroni adjusted). In the Movement group there was no significant main effect of time on RT: F_(5,55)_ = 0.89 p=0.496, partial *η*^2^ = 0.075.

**FIGURE 2.**
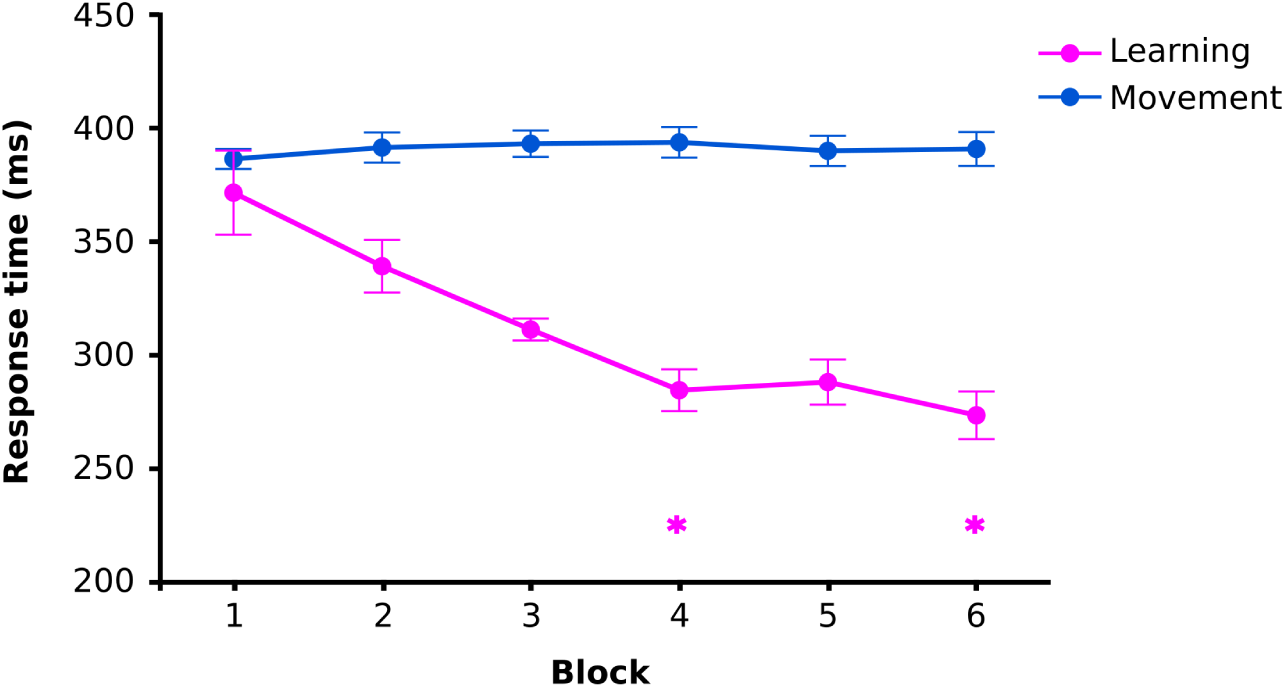
Learning of motor sequence serial reaction time task. Group mean response time data showing a decrease in response times in the Learning SRTT as the participants learned the four-button 16-press sequence (magenta). No equivalent learning was observed in the Movement group SRTT of equivalent duration, which contained no repeating sequence (blue). Two-way Mixed ANOVA: Between-subjects factor: experimental group (Learning or Movement) and Within-subjects factor: time (Block 1-6): F_(1.49,105)_ = 10.52 p=0.001, partial *η*^2^ = 0.334 (Greenhouse-Geisser corrected). This effect was driven by a reduction in response time in the Learning group (Simple main effect of block: F_(1.4,50)_ = 9.33, p=0.005, partial *η*^2^= 0.483, Greenhouse-Geisser corrected). * p<0.05 Bonferroni adjusted post-hoc pairwise comparison compared with block 1. Within-subject error bars calculated across each group (Cousineau, 2005).

In the analysis of MRS GABA:tCr data, a two way mixed ANOVA revealed a significant interaction between experimental group (Learning, Movement, or Rest) and time on GABA:tCr: *F*_(10,155)_ = 2.03, p=0.034, partial *η*^2^= 0.116, (Figure 3) therefore simple main effects were analysed. In the Learning group there was a significant main effect of time on GABA:tCr: *F*_(5,50)_ = 4.16, p=0.003, partial *η*^2^= 0.294. Post-hoc paired-samples t-tests revealed a significant reduction in GABA:tCr in block 6 compared with block 1 (p<0.05, Bonferroni adjusted). There was no significant main effect of time on GABA:tCr in the Movement group (F_(5,55)_ = 1.16 p=0.339, partial *η*^2^ = 0.096), nor in the Rest group (F_(5,50)_ = 0.226 p=0.949, partial *η*^2^ = 0.022).

**FIGURE 3.**
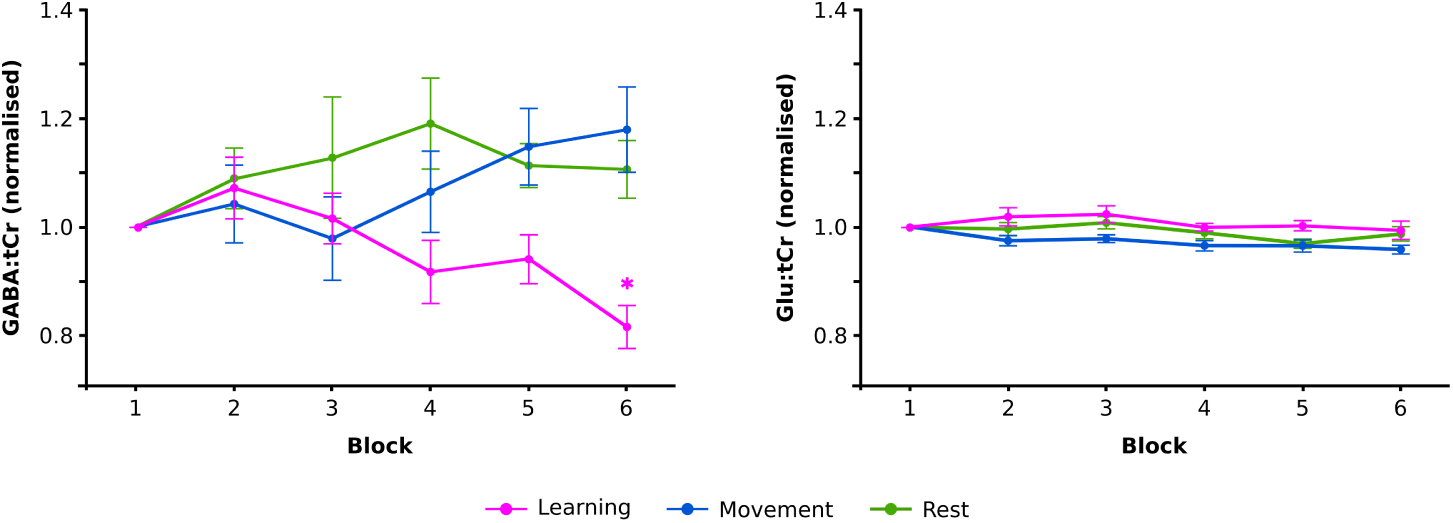
Motor learning is associated with a reduction in motor cortex GABA concentrations. Group mean GABA:tCr and Glu:tCr concentrations presented normalised to block 1 for six serial MRS acquisitions measured during task performance. During motor sequence learning, a reduction in the concentration of motor cortex GABA:tCr is observed (pink) that is not seen in a motor task of equivalent duration lacking a learnable sequence (blue), nor during a passive resting task of the same duration (green). Two-way Mixed ANOVA with one factor of experimental group (Learning, Movement, or Rest) and one factor of block (1-6): *F*_(10,155)_ = 2.03, p=0.034, partial *η*^2^= 0.116. This effect was driven by a drop in GABA:tCr concentration in the Learning group (Simple main effect of block: *F*_(5,50)_ = 4.16, p=0.003, partial *η*^2^= 0.294). Equivalent measures of glutamate showed no evidence of a change specific to the Learning group: a two-way mixed ANOVA revealed no significant interaction between experimental group (Learning, Movement, or Rest) and time on Glu:tCr: F_(10,155)_ = 0.780 p=0.648, partial *η*^2^= 0.048*. p<0.05 Bonferroni adjusted post-hoc pairwise comparison compared with block 1. Within-subject error bars calculated across each group (Cousineau, 2005).

To investigate whether the observed decrease in GABA:tCr during learning might results from longitudinal changes in the fit or SNR of the MRS data specific to the Learning group we performed a two mixed ANOVA on SNR values and CRLB values. These revealed no significant interaction between experimental group (Learning, Movement, or Rest) and time (SNR: p=0.658, CRLB: p=0.273).

### 3.2 Early M1 GABA concentration is predictive of subsequent learning performance

The next question we wished to address was whether levels of M1 inhibition early in learning predicted subsequent learning. This was investigated in the Learning cohort (N=15), including additional participants whose GABA:tCr in block 1 met the CRLB quality criteria (<50%). There was a strong positive correlation between Block 1 GABA:tCr and the peak change in RT, defined as the maximal decrease between RT in block 1 and the RT value of any other block (r(15)=0.658, p=0.0077), such that lower inhibition in M1 is predictive of greater subsequent motor learning (Figure 4A). No such relationship was observed between the baseline measure of M1 Glu:tCr and peak change in RT (r(15)=-0.0408, p=0.8851; Figure 4B). The correlation between M1 GABA:tCr and peak learning was significantly stronger than the equivalent relationship for Glu:tCr (Hittner’s *Z*=2.23, p=0.026). There was no correlation between the magnitude of the change in GABA:tCr and the magnitude of learning (p>0.05).

**FIGURE 4.**
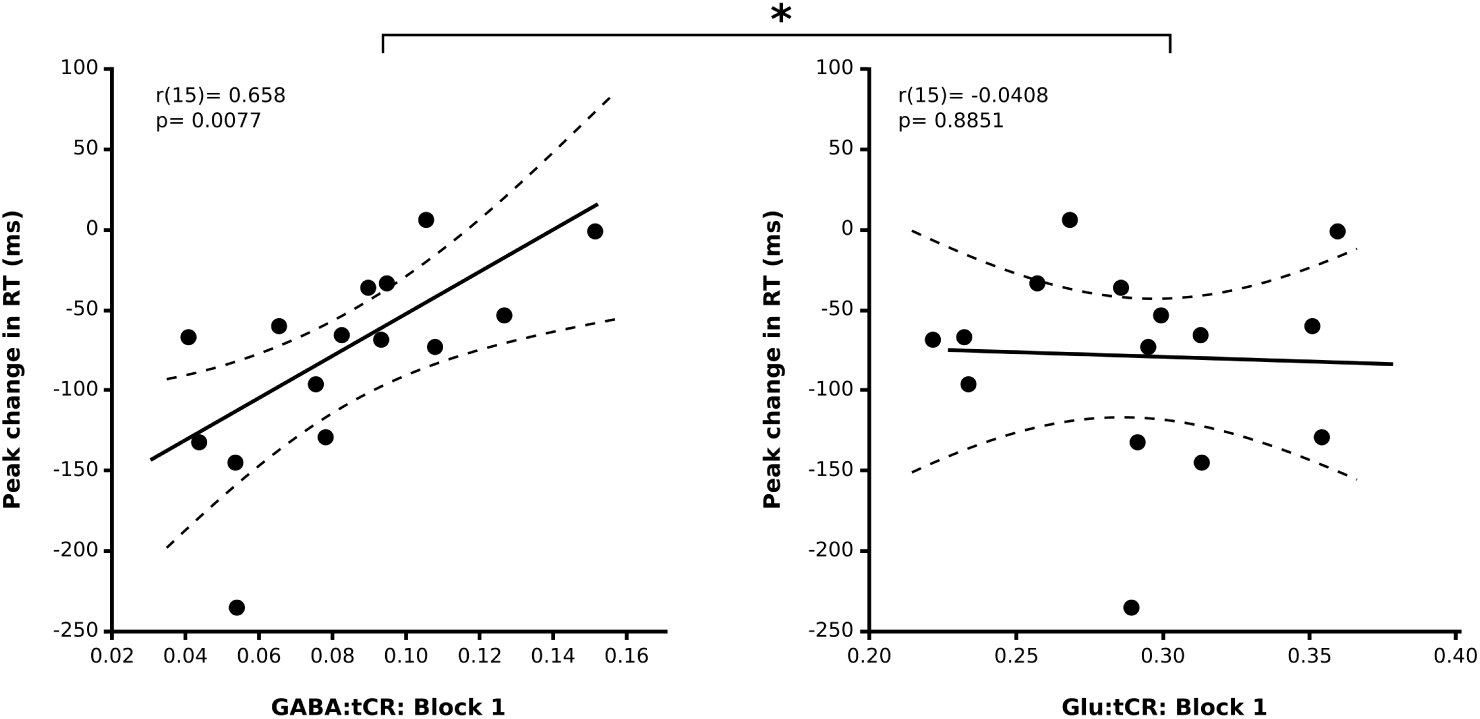
Baseline concentrations of GABA:tCr are strongly correlated with the magnitude of subsequent motor learning. A strong positive correlation was observed between the concentration of GABA:tCr in motor cortex and the peak reduction in response time observed during subsequent motor learning. The same pattern was not observed with equivalent concentrations of excitatory Glutamate (Glu:tCr), suggesting that the level of inhibitory tone present in the sensorimotor cortex may present a barrier to be surmounted for the increase in excitability strongly associated with the acquisition of a new motor skill. *: The correlation of GABA:tCr and Glu:tCr with peak learning differ significantly: Hittner’s *Z*=2.23, p=0.026.

## 4 DISCUSSION

This study was performed to investigate the role of motor cortical GABA in human skill learning. We provide strong evidence for a reduction in the concentration of inhibitory GABA:tCr in M1 early during the acquisition of a learned sequence of movements. The observed reduction in GABA:tCr was specific to the learning observed in the Learning group: it was observed neither in the context of the same finger movements not associated with learning in the Movement group, nor during the Rest group which did not involve finger movements. Furthermore, the observed change in GABA concentration associated with motor learning was not mirrored in changes on glutamate concentrations also derived from MRS. The magnitude of motor learning in the Learning group also strongly predicted by the concentration of baseline GABA:tCr. Taken together these results highlight the potentially crucial and early role for the disinhibition of M1 in supporting a learned movement.

Beyond the phasic GABAergic inhibition central to mechanisms such as lateral inhibition, the tonic activity from extra-cellular GABA is thought to mediate a basal level of inhibitory tone (Semyanov et al., 2004). This ambient inhibitory activity acts via extra-synaptic GABA*_A_* receptors, altering properties such as the membrane refractory period (Glykys and Mody, 2007; Belelli et al., 2009; Isaacson and Scanziani, 2011). This tonic signalling is thought to affect local network activity through a sort of paracrine signalling: the GABAergic overspill from phasic GABA release leads to an tonic extracellular concentration of GABA, whose signalling impacts the excitability of neighbouring neurons (Farrant and Nusser, 2005). MRS measures of GABA concentration are thought to more closely represent the extracellular pool of GABA responsible for this tonic inhibitory activity, rather than the vesicular pool of GABA responsible for phasic signalling, which is bound to macromolecules and therefore less visible to MRS (Stagg et al., 2011b,c). Tonic inhibition, is thought to result from the extrasynaptic overspill of GABA mediating phasic accumulation in the extra-cellular space of GABAergic activity. The evidence of decreased extracellular GABA during learning reported in this study is in keeping with observations of reduced frequency of axonal boutons on inhibitory interneurons immediately after the initiation of training from murine studies of motor learning (Chen et al., 2015): a reduction in the phasic activity of GABAergic neurons may result in a reduction the tonic extracellular GABA pool, which has a subsequent impact on the membrane refractory period and activity of local neurons. The widespread effect of network inhibition is in keeping with observations of heightened M1 excitability after learning (Muellbacher et al., 2001), potentially unmasking latent connections and facilitating plastic change in the connections within M1 (Huntley, 1997).

The strong correlation between the M1 GABA in the earliest stages of task performance and the magnitude of subsequent motor learning (Figure 4 is in keeping with the notion of M1 disinhibition acting as a precursor to the M1 plasticity associated with motor sequence learning. Specifically, greater levels of extracellular GABA acting tonically on local circuits in M1 may prevent or slow the process of local disinhibition associated with learning, such that the magnitude of learning observed in this study was less than that observed in participants whose extracellular GABA concentration was already comparatively low at baseline. In addition, this relationship could also result from the fact individuals showing an early drop in GABA concentration go on to learn more in the study, whereas those who maintain relatively high GABA concentrations early in the task go on to learn less.

We observed no relationship between the magnitude of the reduction in M1 GABA and the change in RT as a measure of learning. However, we are cautious about drawing a firm conclusion from this null finding. It may be that the lack of a relationship between changes in GABA and changes in RT reflects the act that changes in the M1 concentration of GABA reflect only one aspect of learning, which interacts with a variety of other neuronal sub populations and cortical regions (Chen et al., 2015). However, we cannot rule out that the null result could arise from a limitation of the MRS method, particularly the limited temporal resolution of the GABA:tCr measurements. It is also highly likely that the dynamics of the GABA:tCr signal would continue to evolve beyond the 40 minute period of measurement. The time constraint here represents the practical feasibility of acquiring high quality spectra over prolonged continuous periods. Further work focused on understanding the dynamics of the GABA signal during learning could potentially overcome these limitations, and may reveal a relationship between GABA change and learning.

No change in the MRS concentration of glutamate was observed alongside the reduction of GABA in the context of motor learning, movement, or rest. Glutamatergic signalling encompasses a broad range of processes in the cerebral cortex; our results do not exclude the possibility of a change in glutamatergic signalling associated with learning, but rather than our quantification of glutamate may represent a composite of its various roles as both a neurotransmitter and a metabolite. The application of short echo time MRS acquisitions in this study may also have limited the ability to measure changes in the glutamatergic system due to the predominance of a signals from restricted vesicular pools, reducing sensitivity to change (Mullins, 2018). In addition, MRS measures are also not able to quantify changes in glutamate receptor density, which could impact its signalling across the course of learning.

We observed a specific reduction in GABA during Motor Learning; no change in GABA was observed during simple movement. This finding is consistent with the one previous study highlighting reduced MRS-assessed GABA concentrations in human M1 observed during a force-tracking learning task but not in an analogous Movement condition (Floyer-Lea, 2006). However, these results are in contrast to a recent study reporting evidence of a reduction in MRS measures of GABA during a bi-manual whole-hand clench task (Chen et al., 2017). In light of differences in the relative balance of left and right M1 in the context of unimanual versus bimanual tasks (Koeneke et al., 2004), it is difficult to interpret the present results in the context of this study, where the concentration of M1 GABA change may be impacted by a mixture top-down signals and M1-M1 interhemispheric signals that differs from those occurring in a unimanual task.

This work represents an important replication and extension of previous findings regarding the role of M1 inhibition in motor learning. We provide strong evidence for a learning-specific reduction in the measured concentration of M1 GABA, likely to represent a change in the level of local inhibitory tone affected by extrasynaptic GABAergic signalling. Further, we demonstrate a cross-sectional predictive relationship between the concentration of M1 GABA at an early time point in the task and the magnitude of subsequent motor learning, providing initial support for a potential causal link between the set point of local inhibitory tone and the propensity for subsequent plastic change to support behavioural change. Taken together these findings suggest that alterations in inhibitory signalling in M1 likely represent an important step in the mechanism of plasticity that supports motor learning. From a methodological perspective, this paper highlights a step-change in the application of MR spectroscopy, towards more functional-MRS approaches, such that the quantification of trace neurochemicals such as GABA can be tracked over independent scan acquisitions to assess dynamic changes in their concentration in the context of a specific task or exposure.

## ACKNOWLEDGEMENTS

J.K. holds a Wellcome Trust Sir Henry Wellcome Postdoctoral Fellowship (204696/Z/16/Z) and was also supported by a Stevenson Junior Research Fellowship at University College (Oxford) during this work. C.J.S holds a Wellcome Trust/Royal Society Sir Henry Dale Fellowship (102584/Z/13/Z). ELH was additionally supported by the NIHR Oxford Biomedical Research Centre. Support for the 7-T scanner was provided by the Medical Research Council. The Wellcome Centre for Integrative Neuroimaging is supported by core funding from the Wellcome Trust (203139/Z/16/Z).

## SUPPORTING INFORMATION: FOR REVIEW PURPOSES ONLY

**FIGURE 5.**
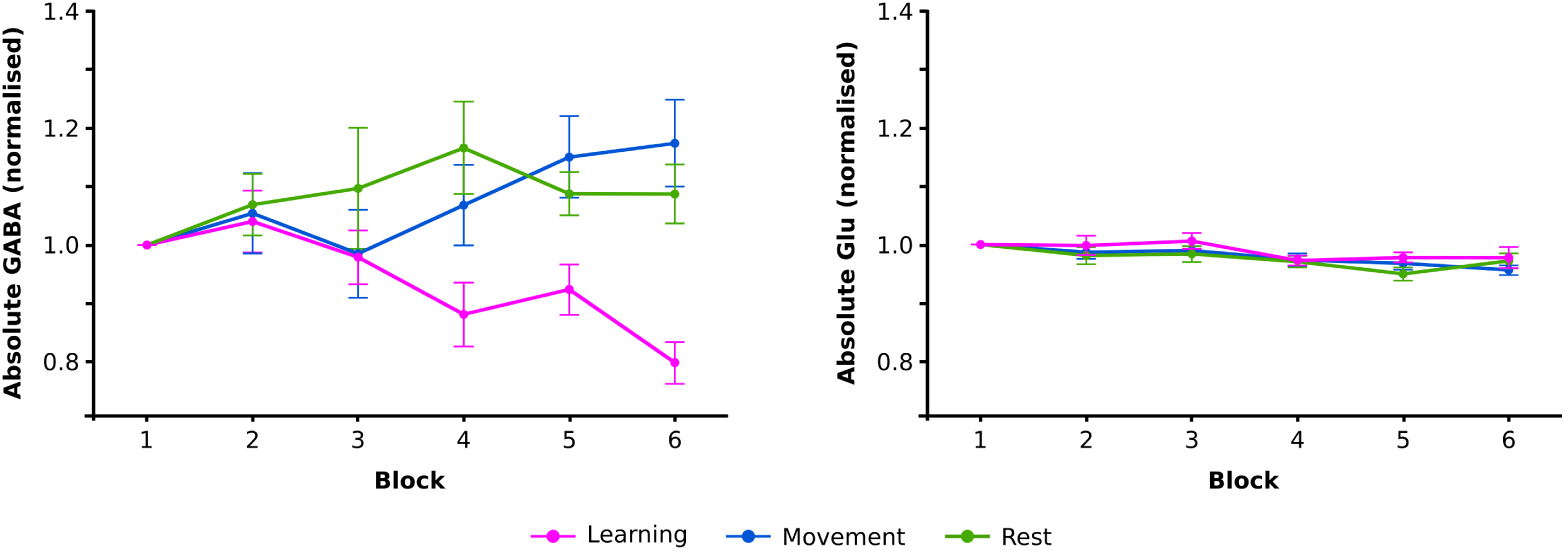
Motor learning associated changes in absolute GABA concentration not corrected for voxel grey matter partial volume or referenced to tCr. Group mean absolute GABA and absolute Glu concentrations presented normalised to block 1 for six serial MRS acquisitions measured during task performance. During motor sequence learning, a reduction in the concentration of motor cortex absolute GABA is observed (pink) that is not seen in a motor task of equivalent duration lacking a learnable sequence (blue), nor during a passive rest task of the same duration (green). Two-way Mixed ANOVA with one factor of group (Learning, Movement, or Rest) and one factor of block (1-6): *F*_(10,155)_ = 2.21, p=0.020, partial *η*^2^= 0.125. This effect was driven by a drop in absolute GABA concentration in the Motor sequence group (Simple main effect of block: *F*_(5,50)_ = 4.25, p=0.003, partial *η*^2^ = 0.298). Equivalent measures of glutamate showed no evidence of a change specific to the Learning group: a two-way mixed ANOVA revealed no significant interaction between experimental group (Learning, Movement, or Rest) and time on absolute glutamate: F_(10,104.70)_ = 0.306 p=0.979, partial *η*^2^= 0.019, Greenhouse-Geisser corrected. * p<0.05 Bonferroni adjusted post-hoc pairwise comparison compared with block 1. Within-subject error bars calculated across each group (Cousineau, 2005).

